# At matched loads, aging does not alter ankle, muscle, or tendon stiffness

**DOI:** 10.1101/2023.11.25.568676

**Authors:** Kristen L. Jakubowski, Daniel Ludvig, Sabrina S.M. Lee, Eric J. Perreault

**Author notes:** Corresponding author Kristen L. Jakubowski.

## Abstract

Older adults have difficulty maintaining balance when faced with postural disturbances, a task that is influenced by the stiffness of the triceps surae and Achilles tendon. Age-related changes in Achilles tendon stiffness have been reported at matched levels of effort, but measures typically have not been made at matched loads, which is important due to age-dependent changes in strength. Moreover, age-dependent changes in muscle stiffness have yet to be tested. Here, we investigate how age alters muscle and tendon stiffness and their influence on ankle stiffness. We hypothesized that age-related changes in muscle and tendon contribute to reduced ankle stiffness in older adults and evaluated this hypothesis when either load or effort were matched. We used B-mode ultrasound with joint-level perturbations to quantify ankle, muscle, and tendon stiffness across a range of loads and efforts in seventeen healthy younger and older adults. At matched loads, there was no significant difference in ankle, muscle, or tendon stiffness between groups (all p>0.13). However, at matched effort, older adults exhibited a significant decrease in ankle (27%; p=0.008), muscle (37%; p=0.02), and tendon stiffness (22%; p=0.03) at 30% of maximum effort. This is consistent with our finding that older adults were 36% weaker than younger adults in plantarflexion (p=0.004). Together these results indicate that, at the loads tested in this study, there are no age-dependent changes in the mechanical properties of muscle or tendon, only differences in strength that result in altered ankle, muscle, and tendon stiffness at matched levels of effort.

**New and Noteworthy:** We provide the first simultaneous estimates of ankle, muscle, and tendon stiffness in younger and older adults. In contrast to earlier conclusions, we found that muscle and tendon mechanical properties are unaffected by age when compared at matched loads. However, due to age-related decreases in strength, mechanical properties do differ at matched efforts. As such, it is important to assess the relevance of the comparisons being made relative to the functional tasks under consideration.

## Introduction

Mobility-related impairments are the leading cause of disability in older adults, affecting nearly 27% of individuals over 65 years old (35). The ability to withstand unexpected postural disturbances is critical for healthy mobility and fall prevention (20, 22). Older adults have a reduced capacity to compensate for postural disturbances (39), which increases their risk of falling (26, 46). For example, one-third of falls in older adults are directly related to the inability to respond appropriately to postural disturbances (12). An appropriate response requires rapid corrective actions that oppose the disturbance. The ankle is crucial in this response, producing a substantial portion of the required torque (41). This torque arises from both the stiffness of the ankle at the time of the perturbation and from delayed muscle activation mediated by sensory feedback pathways (11, 15, 16). Previous investigations of older adults’ impaired response to postural disturbances have emphasized the role of age-dependent changes in the sensory feedback pathways (1, 26, 47), but there has been limited investigation into how aging impacts ankle stiffness. If ankle stiffness decreases, the importance of sensory-mediated feedback responses would be increased. This is important not only for understanding age-dependent changes in the neural control of posture and balance, but also for designing interventions that could reduce falls since training targeting deficits in neural feedback will differ from that targeting biomechanical deficits in ankle stiffness.

Age-related changes within the triceps surae and Achilles tendon may decrease ankle stiffness in older adults. The triceps surae and Achilles tendon are the primary contributors to sagittal plane ankle stiffness during standing, with ankle stiffness being very sensitive to the stiffness of the Achilles tendon at higher loads (18). It has been suggested that older adults exhibit a decrease in Achilles tendon stiffness (4, 27, 44), which may be associated with impaired mobility and a decreased ability to respond to postural disturbances (7, 10). However, it is worth highlighting that due to methodological assumptions, nearly all previous estimates have been made at higher loads (above 30% MVC) that are less relevant to everyday tasks like standing and walking, where plantarflexor muscle activation typically remains below 30% MVC (33). To our knowledge, it has yet to be explored if there are age-related changes in tendon stiffness at the lower load regime within the "toe-region" of the tendon’s non-linear stress-strain relationship. Additionally, nearly all previous examinations into age-related differences in Achilles tendon stiffness have been made at matched efforts (27, 44). If the goal is to understand differences in mechanical properties, older adults’ decrease in strength presents a confounding factor. In contrast to the Achilles tendon, age-related changes in muscle stiffness remain poorly characterized. To our knowledge, measures of muscle stiffness in younger and older adults during active conditions relevant to postural control have not been made. Despite the lack of experimental measures, age-related changes—such as a progressive decrease in muscle size, an increase in connective tissue, and an increase in fatty infiltration (5, 32, 44)—could impact muscle stiffness. Collectively, these age-related changes within the muscle and tendon could alter ankle stiffness in older adults.

Our primary objective was to determine if ankle, muscle, and tendon stiffness differ between younger and older adults. We used our measurement technique that combines B-mode ultrasound imaging with joint-level perturbations to quantify ankle, muscle, and tendon stiffness simultaneously (17). Our approach allowed us to overcome gaps in the literature so that muscle and tendon stiffness could be estimated at lower activation levels — from 0 to 30% MVC. Based on the previously reported age-related decrease in Achilles tendon stiffness (4, 27, 44), we hypothesize that older adults will exhibit a decrease in Achilles tendon stiffness that will result in a decrease in ankle stiffness at matched levels of load and effort. Our results provide new insight into the mechanisms underlying age-dependent differences in the ability to regulate ankle mechanics.

## Methods

### Participants

Seventeen healthy younger adults and seventeen healthy older adults participated in this study (Table 1). All participants were right-leg dominant, as determined by the revised Waterloo Footedness Questionnaire (9). Participants completed a health questionnaire prior to participation to ensure they had no history of musculoskeletal injury or surgery to their right leg, neurological diseases or injuries, or were taking medication that causes dizziness or impacts balance. Older adults completed two additional questionnaires that evaluate fall risk and activity level: the Center for Disease Control (CDC) fall risk self-assessment (Stay Independent Brochure) (40), and the Community Healthy Activities Model Program for Seniors (CHAMPS) (45). The fall risk self-assessment asked 12 yes-no questions. The "yes" are summed, and a score above 4 indicates the individual may be at a higher risk of falling. A higher score on the CHAMPS questionnaire indicates the individual is more active. All participants provided informed consent before participating in the study. The Northwestern University Institutional Review Board (IRB) approved the study, and all methods were carried out in accordance with the IRB-approved protocol (STU00009204 & STU00213839).

**Table 1:**
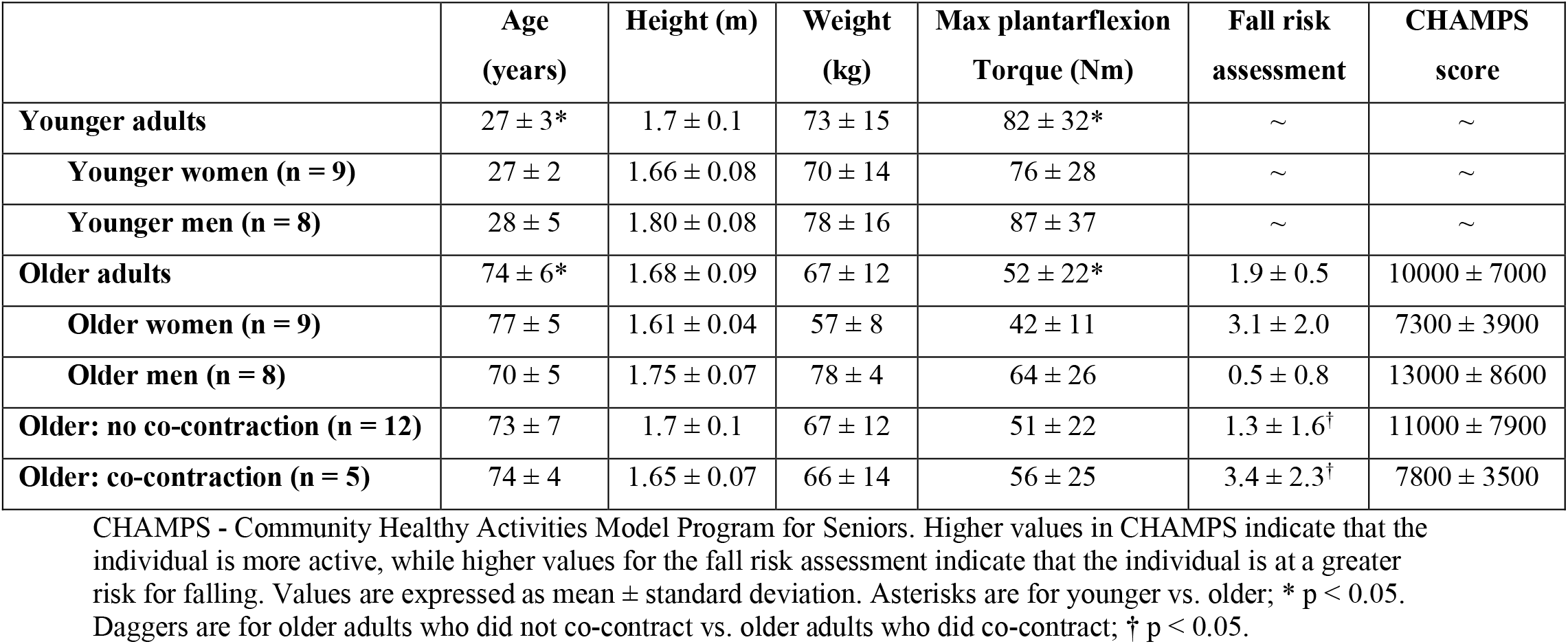
Subject characteristics.

### Experimental setup

Participants sat in an adjustable Biodex chair (Biodex Medical Systems, Inc. Shirley, NY) with their right leg extended in front of them (Fig 1A). Their knee was flexed at 15° and stabilized with a brace (Innovator DLX, Ossur, Reykjavik, Iceland). The torso and trunk were stabilized with safety straps. The right foot was rigidly secured to a brushless rotary motor (BSM90N-3150AF, Baldor, Fort Smith, AR) via a custom-made fiberglass cast with the ankle positioned at 90° of flexion. The cast encased the foot distally from the malleoli to beyond the toes, creating a rigid foot while preserving the full range-of-motion of the ankle. The ankle center of rotation in the sagittal plane was aligned with the axis of rotation of the motor, restricting all movement to the plantarflexion/dorsiflexion direction. Electrical and mechanical safety stops limited the rotation of the motor within the participant’s range of motion. A 24-bit quadrature encoder integrated with the motor measured ankle angle (24-bit, PCI-QUAD04, Measurement Computing, Norton, MA), while a 6-degree-of-freedom load cell (45E15A4, JR3, Woodland, CA) measured all ankle forces and torques. The motor was controlled in real-time via xPC target throughout the experiment (MATLAB, Mathworks, Natick, MA).

**Figure 1.**
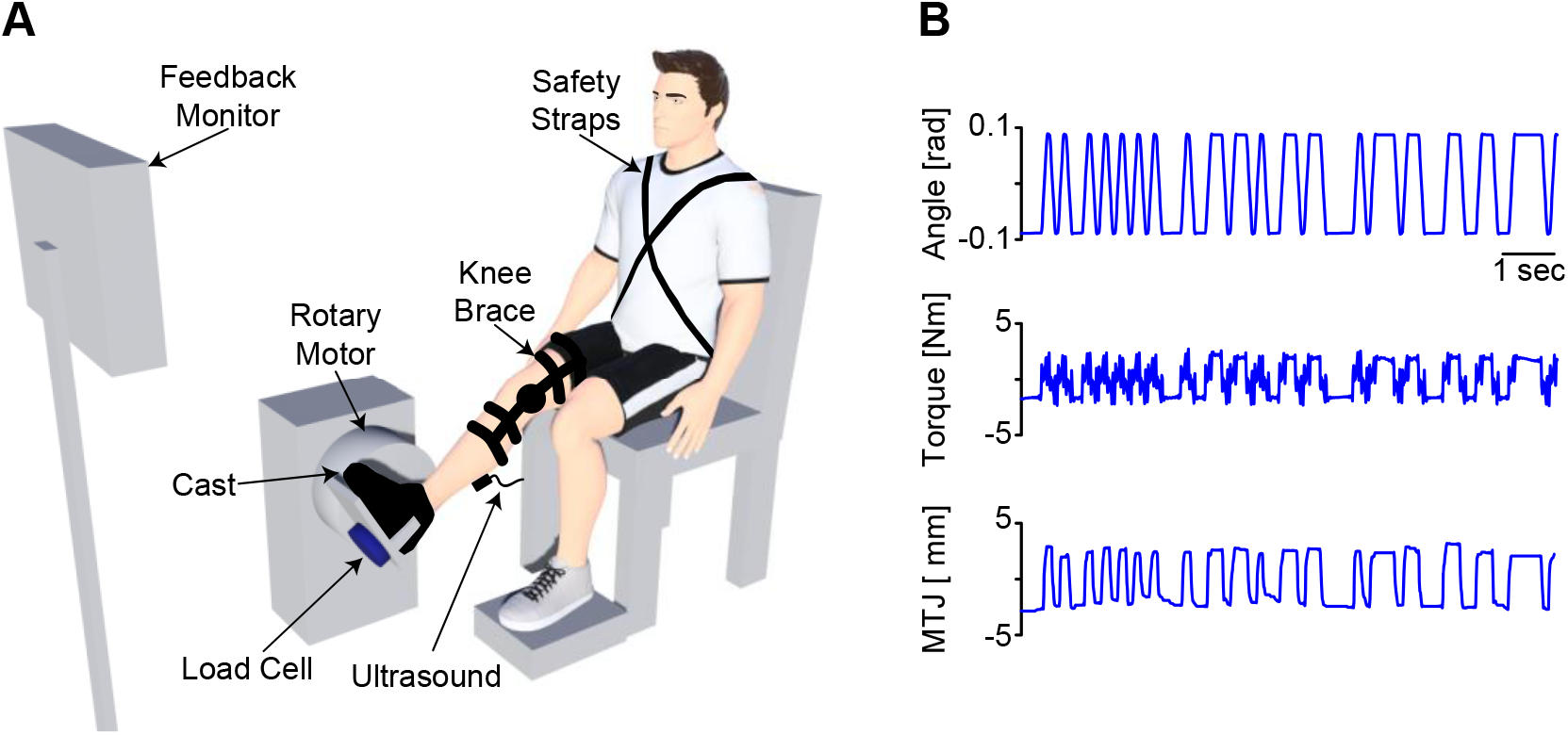
(A) Schematic of the experimental setup. (B) Representative data used to estimate ankle, muscle, and tendon stiffness for an older adult. A custom-made cast secured the participant’s foot to the rotary motor. The rotary motor rigidly controlled the ankle joint angle and applied random perturbations while the load cell measured the resultant ankle torque. We used B-mode ultrasound to image the muscle-tendon junction (MTJ) of the medial gastrocnemius. The knee brace secured the knee in a stable position, preventing unwanted knee flexion or extension. The feedback monitor provided real-time feedback on the magnitude of the plantarflexion torque and the tibialis anterior muscle activity. Figure adapted from Jakubowski et al. (18). (B) Exemplary measures of the ankle angle, torque, and MTJ displacement resulting from the applied random perturbations.

Electromyography (EMG) data from the medial and lateral gastrocnemius and soleus (ankle plantarflexors) and the tibialis anterior (ankle dorsiflexor) were recorded using single differential bipolar surface electrodes (Bagnoli, Delsys Inc, Boston, MA, 10 mm interelectrode distance). We performed standard skin preparation techniques prior to placing each electrode on the belly of the respective muscle (48). All analog data were passed through an antialiasing filter (500-Hz 5-pole low-pass Bessel filter) and sampled at 2.5 kHz (PCI-DAS1602/16, Measurement Computing, Norton, MA, USA). EMG data were collected and used to provide visual feedback to the participant.

We collected B-mode ultrasound images of the medial gastrocnemius muscle-tendon junction (MTJ) using a linear transducer (LV7.5/60/128Z-2, LS128, CExt, Telemed, Lithuania). The MTJ was positioned in the center of the ultrasound image. The ultrasound probe was secured to the leg using a custom-made probe holder and elastic adhesive wrap (Coban™, 3M, St. Paul, MN). A trigger synchronized the ultrasound data with all other measurements. Ultrasound images were acquired with a mean frame rate of 124 frames per second. All ultrasound data were saved for processing offline.

### Protocol

Participants completed isometric maximum voluntary contractions (MVC) at the start of the experiment. These data were used to normalize the EMGs and scale the torque and EMGs for visual feedback provided to the participants during later trials (2). Participants completed three 10-second MVC trials in both plantarflexion and dorsiflexion directions.

Our primary objective was to determine how the triceps surae, Achilles tendon, and ankle impedance varied between younger and older adults across different levels of plantarflexion torque. This contrasts previous studies that evaluated age-related changes in Achilles tendon stiffness at matched efforts (27, 44). Therefore, the rotary motor applied small rotational perturbations in the sagittal plane while participants produced different levels of isometric plantarflexion torque. We used pseudo-random binary sequence (PRBS) perturbations with an amplitude of 0.175 radians, a maximum velocity of 1.75 radians per second, and a switching time of 153 ms. We tested seven plantarflexion torque levels from 0% to 30% MVC in 5% increments, with participants completing three trials at each level of plantarflexion torque. Participants were provided real-time visual feedback of their normalized plantarflexion torque along with tibialis anterior EMG. Subjects were instructed to match the target torque for each trial while minimizing tibialis anterior EMG to avoid co-contraction. Rectified EMG and torque signals used for the visual feedback were low pass filtered at 1 Hz to remove high-frequency components (2^nd^-order Butterworth). Each trial lasted 65 seconds. Plantarflexion torque levels were tested in a randomized fashion. Rest breaks were provided as needed between trials to prevent fatigue.

The measured ankle torque included the gravitational and inertial contributions from the apparatus connecting the foot to the motor. Thus, we collected a single trial with only the cast attached to the rotary motor. This enabled us to remove the contributions from the apparatus from the net torque measured in each trial.

### Data processing and analysis

We processed and analyzed all data using custom-written software in MATLAB. The same experimenter manually digitized the MTJ within each frame of all ultrasound videos (17). All ultrasound metrics were synchronized with all other data (29) and resampled using linear interpolation to match the sampling rate of all other data (2.5 kHz).

We examined the EMG data to determine if there were age-related differences in muscle activation. All EMG data were notch-filtered to remove 60 Hz noise, detrended, and rectified. The EMG signals were smoothed with a 25-ms moving average filter, and the average across the trial was taken. The EMG signals were then normalized by the peak amplitude of the filtered EMG signal from the MVC trials.

We used non-parametric system identification to compute ankle, muscle, and tendon impedance, as previously described (17). Briefly, the experimental measures used in this analysis were ankle angle, ankle torque, and MTJ displacement (Fig 1B). Ankle impedance was quantified as the relationship between the imposed ankle rotations and the resultant ankle torque (19). We modeled the triceps surae and Achilles tendon as two impedances in series (14), where the displacement of the muscle-tendon unit is determined by the angular rotation of the ankle multiplied by the Achilles tendon moment arm. We refer to the relationship between MTJ displacement and the angular rotations of the ankle as the translation ratio. Specifically, to characterize ankle, muscle, and tendon impedance, we estimated ankle impedance and the translation ratio. We then used these quantities to compute muscle and tendon impedance algebraically. We have previously demonstrated that the magnitude of the frequency response functions were nearly constant from 1 to 3 Hz and had high coherence, indicating that stiffness is the dominant contributor to impedance and that there is a high signal-to-noise ratio (17). Thus, we computed the stiffness component of ankle, muscle, and tendon impedance by averaging the magnitude of the frequency response functions from 1 to 3 Hz. Our analysis focuses on the stiffness component of impedance due to its relevance in the control of posture and movement at the ankle (24).

For all analyses, we used a single approximation of the Achilles tendon moment arm, taken as the mean across subjects from Clarke et al. (3), with an ankle angle of 90 degrees (51.4mm). A single approximation was deemed appropriate since system identification is a quasi-linear approximation about a single operating point, which, in our study, was 90°. Moreover, it has been demonstrated that the Achilles tendon moment arm does not scale with anthropometric data (3, 42), nor is it affected by age (8, 28).

Ankle and tendon stiffness, the low-frequency component of impedance, varied non-linearly with plantarflexion torque (or musculotendon force). Therefore, the ankle and tendon stiffness experimental data were fit with non-linear mixed-effects models for further analysis, as described previously (18). Torque-dependent changes in ankle stiffness were modeled by:

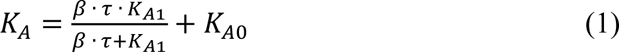

where *K_A_* represents the modeled ankle stiffness, torque (*τ*) was the input to the model, and *β*, *K_Α1_*, and *Κ_Α0_* were the optimized parameters. *K_A0_* represents the passive stiffness of the ankle. A similar approach, excluding *K_A0_*, has been used to characterize the load-dependent changes in net musculotendon stiffness (31).

An exponential function was used to model tendon stiffness:

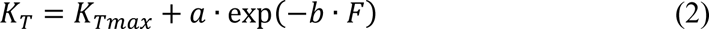

where *K_T_* represents the modeled tendon stiffness, musculotendon force (*F*) was the input to the model, and *K_Tmax_*, *a*, and *b* were the optimized parameters. This model was chosen since exponential models have been used previously to model the non-linear toe-region of the tendon stress-strain curve (23). We computed musculotendon force by dividing the measured ankle torque by the Achilles tendon moment arm. We want to note that this non-linear model of tendon properties (Eq 2) was not incorporated into Eq 1). We made this simplification for two reasons; first, Eq 1 has been previously used in the literature and fit our data well (6, 18, 31). Second, we were unable to fit a model of ankle stiffness that included the non-linear mechanics of the Achilles tendon due to high parameter covariance that led to poor convergence.

### Statistical analysis

We tested the hypothesis that ankle, muscle, and tendon stiffness differed between younger and older adults. Non-linear mixed-effects models were used to characterize ankle and tendon stiffness (Eq 1 and 2), whereas a linear mixed-effects model was sufficient to describe the relationship between muscle stiffness and force. We also examined how ankle, muscle, and tendon stiffness varied with effort (% MVC). Ankle, muscle, and tendon stiffness all varied linearly with % MVC; thus, we used linear mixed-effects models. For all models, age group (younger and older) was treated as a nominal fixed factor, plantarflexion torque, musculotendon force, or % MVC was a continuous factor, and subject was a random factor. We tested our hypothesis by evaluating how age group (older vs. younger) influenced stiffness at torques (or musculotendon forces) ranging from zero to the 75^th^ quantile of the maximum measured torque (or musculotendon force) in older adults, or from efforts ranging from 0 to 30% MVC. We performed a post-hoc analysis to determine if stiffness (ankle, muscle, and tendon) significantly differed between younger and older adults at matched loads (torques or musculotendon forces) or matched efforts (% MVC). We assessed the fit of each model by quantifying the coefficient of determination (R^2^) for each participant from the respective mixed-effects model. For all mixed-effects models used in our analysis, we used a restricted maximum likelihood method when approximating the likelihood of the model, and Satterthwaite corrections for degrees of freedom (25). We used a two-sample *t-test* to determine if there was a significant difference in maximum plantarflexion torque, *K_Tmax_* (from Eq 2), or subject characteristics between younger and older adults. We performed all statistical analyses in MATLAB. Significance was set *a priori* at α=0.05. All metrics are reported as the mean±95% confidence intervals unless otherwise noted.

## Results

### Participant characteristics

As a group, older adults (n=17) were 36% weaker in their maximum plantarflexion torque compared to younger adults (n=17; p=0.004; Table 1). There was no significant difference in height and weight between younger and older adults (all p>0.15; Table 1). Five older adults were classified as co-contracting (details below). This group scored significantly higher on the fall risk assessment than the eleven older adults who did not co-contract (p=0.04; Table 1). No other differences were observed between the older adults who co-contracted and those who did not (all p>0.5; Table 1).

### Aging did not alter ankle stiffness

Ankle stiffness increased with plantarflexion torque in all younger and older adults; however, no age-related differences were observed (Fig 2). Figure 2A displays the experimental measures and the fits to Eq 1 for representative younger and older adults. The model fit the data well for both groups of participants (younger: R^2^=0.98; older: R^2^=0.99). Similar fit accuracies were achieved across the entire cohort (younger: R^2^=0.98 ± 0.01; older: R^2^=0.96 ± 0.02). We compared the modeled ankle stiffness for younger and older adults at matched torques and found no statistical differences within the range of torques considered in this study (Fig 2B & C). At the 75^th^ quantile of torque measured in the older adult population (∼21 Nm), ankle stiffness was 11% lower in older adults (younger: 134 ± 6 Nm/rad; older: 119 ± 17 Nm/rad; p=0.19). While there was no significant difference in ankle stiffness between the populations in the range of loads tested here, there was more variability in the estimated stiffness from older adults, as shown by the larger confidence bounds in Fig 2B. We next explored if this higher variability was associated with inconsistent levels of co-contraction.

**Figure 2.**
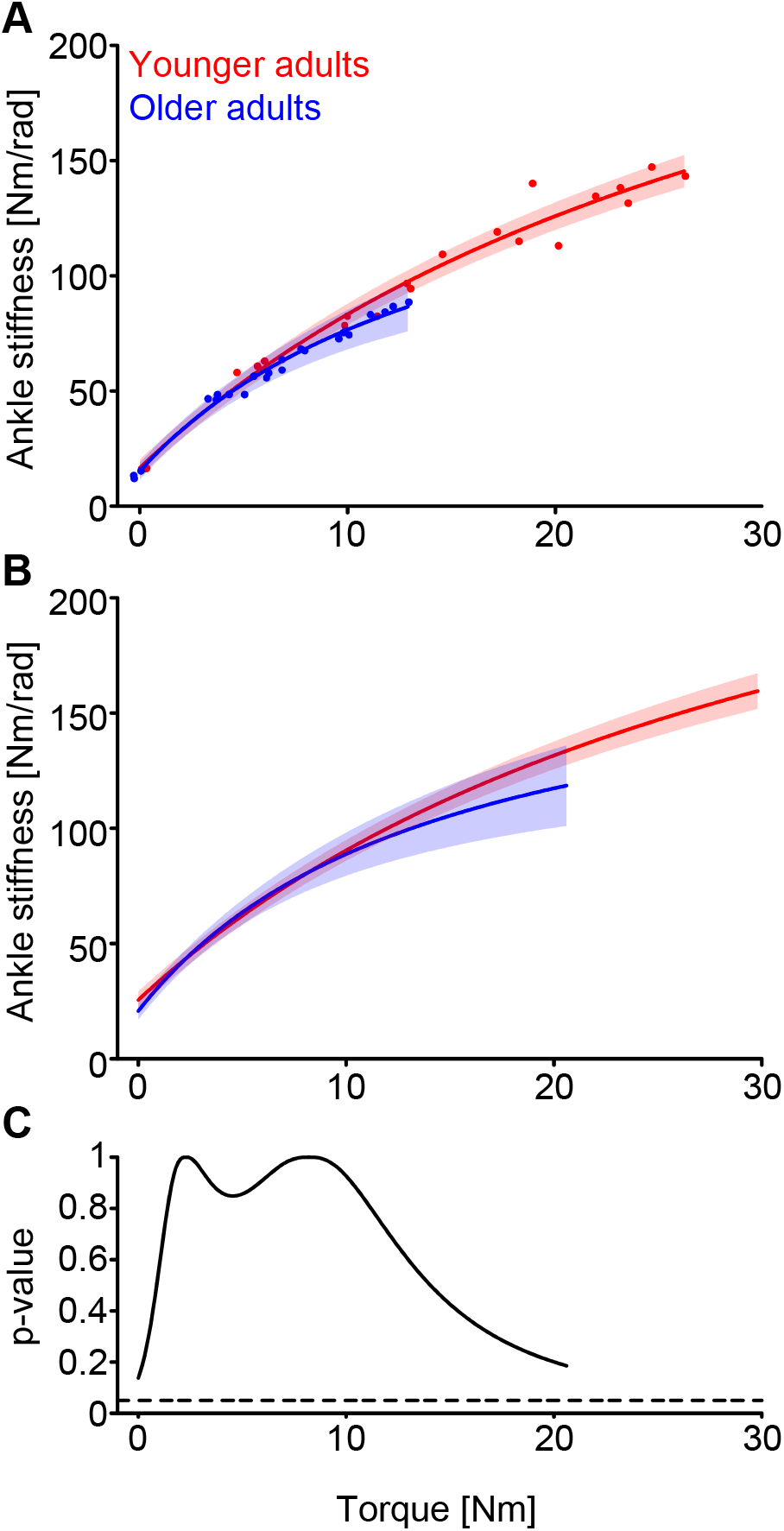
Ankle stiffness was not significantly different in older adults compared to younger adults. (A) Ankle stiffness for a representative younger (red) and older (blue) adult. Younger adults were stronger than older adults, as illustrated by the lower torque achieved in the representative older adult. (B) This trend was preserved in the group results for younger (n=17) and older (n=17) adults. Ankle stiffness data were modeled using a non-linear mixed-effects model (Eq 1). The mixed-effects model was used to account for random variability between participants. The solid line indicates the estimated stiffness from the fitted model, with the shaded region being the 95% confidence intervals. (C) Ankle stiffness was not significantly different between younger and older adults at matched levels of plantarflexion torque, illustrated by the p-value being greater than 0.05 (dotted line). Comparisons were made at torques ranging from zero to the 75^th^ quantile of the maximum measured torque in older adults.

### Aging increases co-contraction about the ankle

On average, older adults exhibited significantly higher tibialis anterior activation compared with younger adults (difference: 2.4 ± 0.9% MVC; p=0.003; Fig 3), indicative of co-contraction in the plantarflexion task used in our protocol. This occurred even though visual feedback of tibialis anterior activation was provided to minimize co-contraction. The increase in tibialis anterior activation contributed to changes in ankle stiffness at matched levels of plantarflexion torque. Figure 4 shows a representative older adult who exhibited co-contraction during some but not all trials. At nearly matched levels of plantarflexion torque (∼ 4 Nm; Fig 4 – black box), ankle stiffness was 64 ± 4 Nm/rad (mean ± standard deviation) at low levels of co-contraction, marked by tibialis anterior activity of 2.1 ± 0.9% MVC (mean ± standard deviation; Fig 4 – filled circles). Stiffness increased to 77 ± 2 Nm/rad as co-contraction increased (tibialis anterior: 6 ± 2% MVC; mean ± standard deviation; Fig 4 – open circles). These results indicate that the increased co-contraction observed in older adults may have contributed to their increased variability in ankle stiffness.

**Figure 3.**
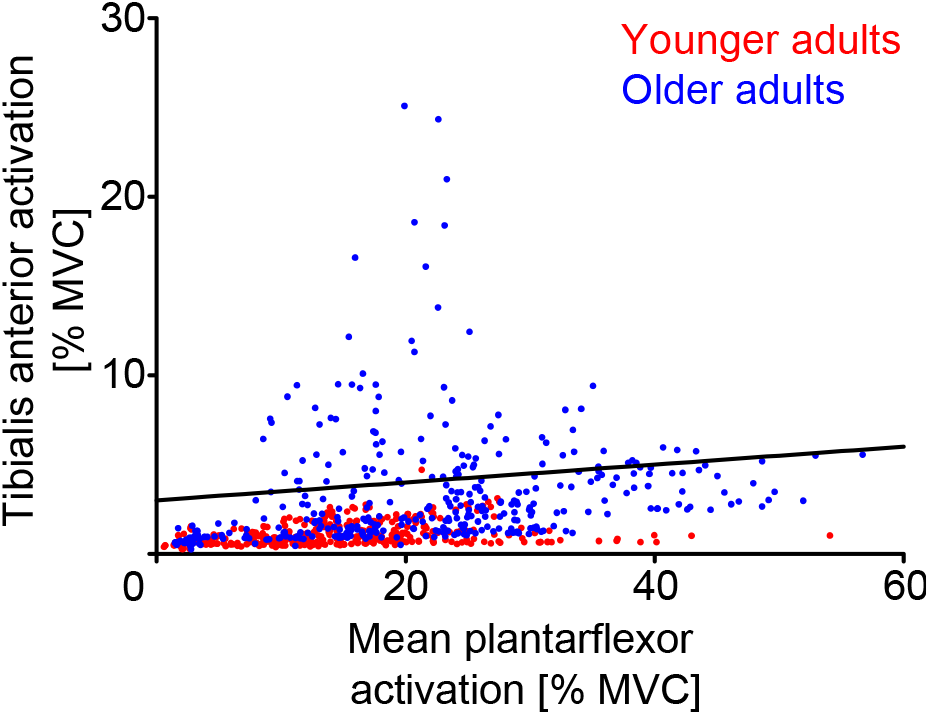
Older adults co-contracted more than younger adults. As the mean plantarflexor activation increased, there was a greater increase in tibialis anterior activation in older adults (n = 17) compared to younger adults (n = 17). Mean plantarflexor activation was computed as the mean activation within the lateral gastrocnemius, medial gastrocnemius, and soleus. Co-contraction was defined as tibialis anterior activation exceeding 5% of the mean plantarflexor activity with a 3% offset (black line). Each data point represents an individual trial in younger (red) and older adults (blue).

**Figure 4.**
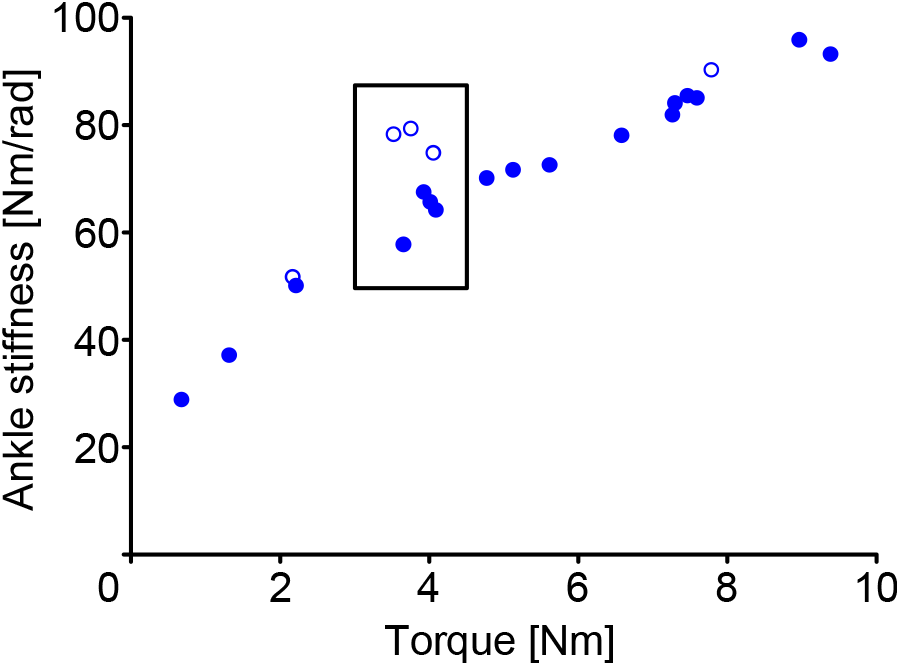
A representative older adult that exhibited co-contraction during some trials. At nearly matched levels of plantarflexion torque (black box), ankle stiffness was higher when the participant co-contracted (open circles) versus when co-contraction was not present (filled circles). Co-contraction was defined as tibialis anterior activation exceeding 5% of the mean plantarflexor activity with a 3% offset. Each data point represents an individual trial.

We removed subjects with significant co-contraction from all subsequent analyses since co-contraction violates an assumption of our method. The assumption is that all perturbation-induced torques (and forces) are transmitted through the Achilles tendon. Subjects were removed if more than 30% of their trials exhibited excessive tibialis anterior activation. Excessive activation was defined as mean tibialis anterior activity exceeding 5% of the mean plantarflexor activity; a 3% offset was used to avoid removing trials at very low torque levels (Fig 3- black line). Twelve of the seventeen older adults and all younger adults were classified as not co-contracting. Within this cohort, individual trials were removed if the tibialis anterior activation exceeded the cut-off described above (0.3% of data for younger adults; 11% of data for older adults). The tibialis anterior activation within this subset of older adults was still significantly higher than younger adults. However, the activations were more similar, and the difference was quite small (0.8 ± 0.4% MVC; p=0.008). We would like to note that we evaluated different criteria for defining co-contraction (e.g., mean tibialis anterior activity above 3% of plantarflexor activity); using different criteria did not influence our main results.

### Aging did not alter muscle or tendon stiffness at matched loads

We evaluated if there were age-related differences in ankle, muscle, and tendon stiffness in the twelve older adults who did not co-contract. Similar to the results presented above, ankle stiffness was not significantly different at matched levels of plantarflexion torque (Fig 5C & F). At the 75^th^ quantile of torque that we measured in older adults (∼21 Nm), ankle stiffness of the older adults was 10% lower than the younger adults (younger: 134 ± 7 Nm/rad; older: 121 ± 13 Nm/rad; p=0.18).

**Figure 5.**
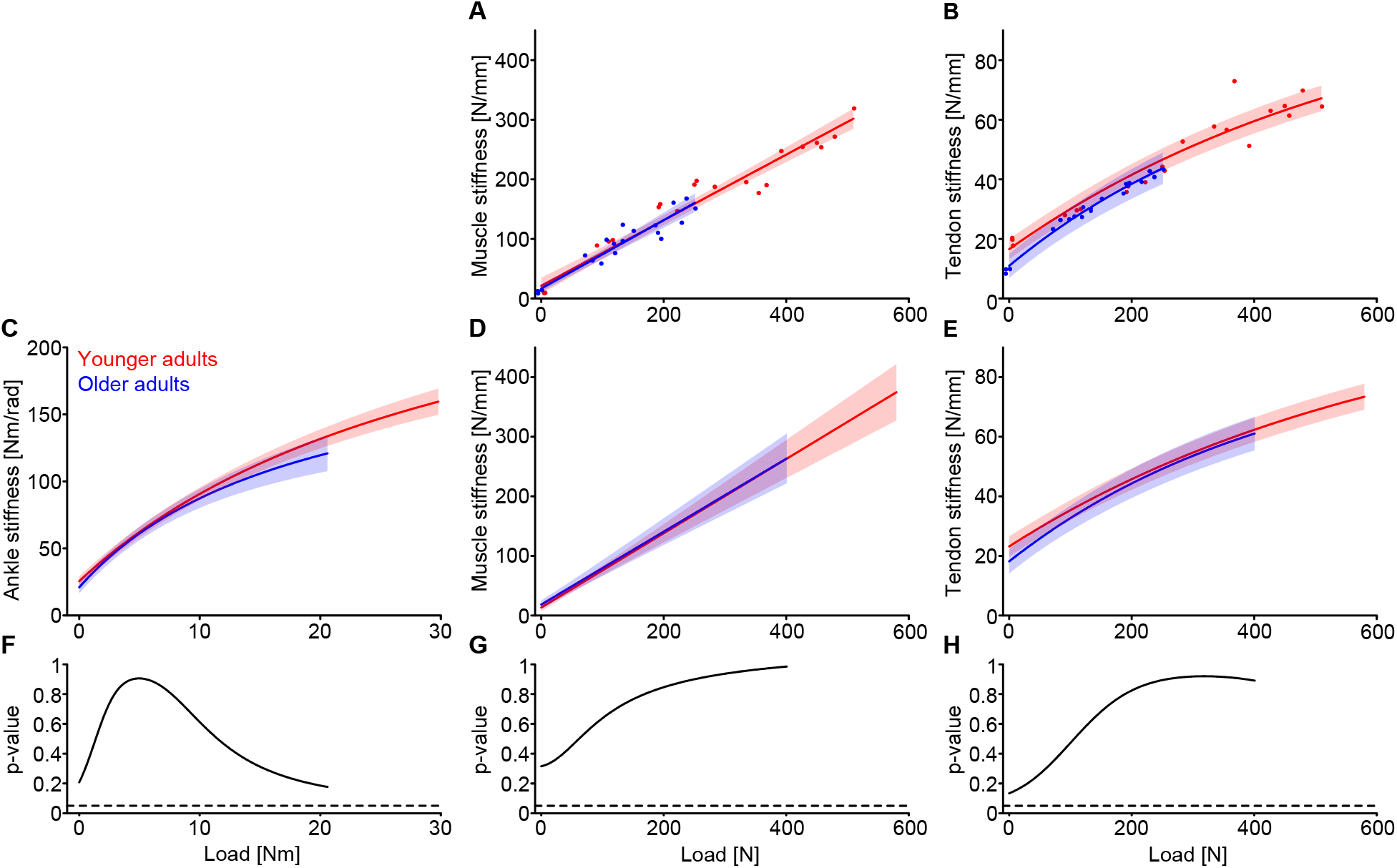
At matched loads, ankle, muscle, and tendon stiffness did not vary between younger and older adults who did not co-contract. Muscle (A) and tendon (B) stiffness for a representative younger (red) and older (blue) adult. (C) Ankle, (D) muscle, and (E) tendon stiffness for younger adults (n=17) and older adults who did not co-contract (n=12). No significant difference was observed in ankle (F), muscle (G), or tendon (H) stiffness between younger and older adults. The ankle stiffness experimental data were modeled using a non-linear mixed-effects model (Eq 1). Similarly, the tendon stiffness experimental data were modeled non-linearly (mixed-effects model) with Eq 2. Muscle stiffness was modeled using a linear mixed-effects model. The solid line indicates the estimated stiffness from the respective fitted model, with the shaded region being the 95% confidence intervals. Comparisons were made at torques ranging from zero to the 75^th^ quantile of the maximum measured torque (or musculotendon force) in older adults.

As musculotendon force increased, muscle stiffness increased linearly, while tendon stiffness increased non-linearly (Fig 5). Figure 5A displays the experimental measures and model fits for muscle stiffness for a representative younger and older adult. The model fit the data well for both participants (younger: R^2^=0.94; older: R^2^=0.89), and across the entire cohort (younger: R^2^=0.95 ± 0.01; older: R^2^=0.93 ± 0.02). Similarly, Figure 5B displays the experimental measures and the parametric model (Eq 2) of the relationship between tendon stiffness and force in a representative younger and older adult. The model fit the data well for both participants (younger: R^2^=0.92; older: R^2^=0.99), and across the entire group (younger: R^2^=0.94 ± 0.02; older: R^2^=0.94 ± 0.04).

At matched levels of musculotendon force, muscle stiffness (Fig 5G) and tendon stiffness (Fig 5H) were not significantly different between younger and older adults. We compared stiffnesses at the 75^th^ quantile of musculotendon force measured in older adults (∼400 N) and found no significant differences in muscle (younger: 263 ± 32 N/mm; older: 263 ± 42 N/mm; p=0.99) or tendon stiffness (younger: 62 ± 4 N/mm; older: 61 ± 6 N/mm; p=0.89) between groups.

Nearly all previous estimates of tendon stiffness have been made at a matched effort (e.g., % MVC) rather than at a matched load (e.g., torque or musculotendon force). Thus, we also examined how ankle, muscle, and tendon stiffness varied between younger and older adults at matched efforts (Fig 6). Figure 6A, B, and C display the experimental measures and the model fits from a representative younger and older adult. Again, the models fit the data well, with high R^2^ across the entire group (*Ankle*: younger: R^2^=0.96 ± 0.01; older: R^2^=0.95 ± 0.02; *Muscle*: younger: R^2^=0.95 ± 0.01; older: R^2^=0.93 ± 0.02; *Tendon*: younger: R^2^=0.93 ± 0.02; older: R^2^=0.91 ± 0.04). At matched effort, ankle, muscle, and tendon stiffness were significantly lower in older adults compared to younger adults. At 30% MVC, the highest target effort in the experiment, ankle stiffness was 27% lower (younger: 153 ± 20 Nm/rad; older: 112 ± 20 Nm/rad; p=0.008, Fig 6G), muscle stiffness was 37% lower (younger: 313 ± 74 N/mm; older: 199 ± 49 N/mm; p=0.02, Fig 6H), and tendon stiffness was 22% lower (younger: 68 ± 8 N/mm; older: 53 ± 9 N/mm; p=0.03, Fig 6I). These differences are driven by differences in strength between the old and young participants.

**Figure 6.**
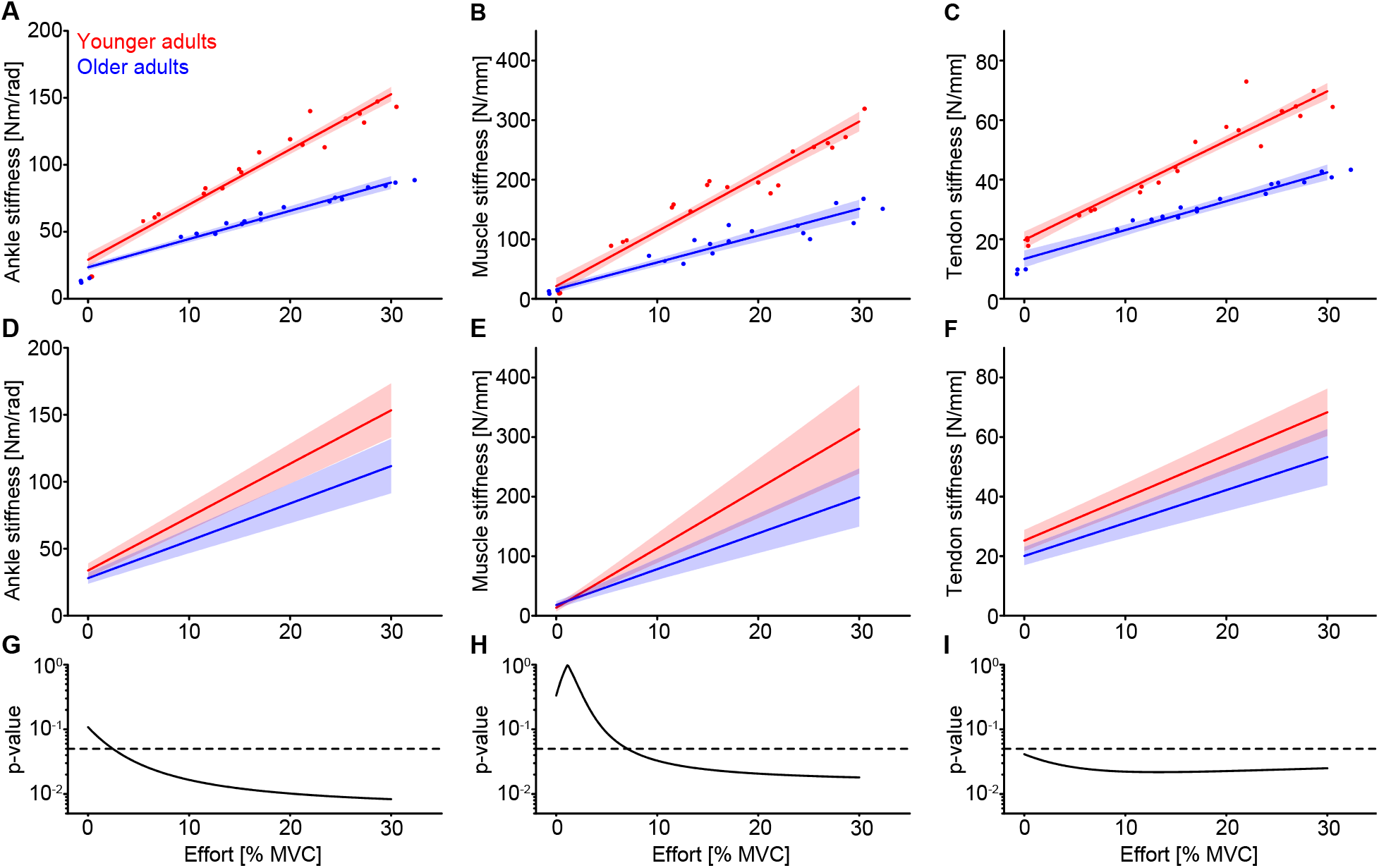
At matched levels of effort, older adults exhibited a decrease in ankle, muscle, and tendon stiffness. Ankle (A), muscle (B), and tendon (C) stiffness for a representative younger (red) and older (blue) adult as a function of effort (% MVC). Ankle, muscle, and tendon stiffness increased linearly with effort, and this trend was preserved when looking at the group results for younger (n=17) and older adults who did not co-contract (n=12) (D, E, and F, for ankle, muscle, and tendon stiffness respectively). At almost all efforts, ankle and tendon stiffness were significantly lower in older adults compared with younger adults (G and I, respectively). Muscle stiffness was significantly lower in older adults compared with younger adults at efforts above approximately 7 % MVC. Note that p-values are plotted on a log scale. Ankle, muscle, and tendon stiffness were modeled using a linear mixed-effects model. The solid line indicates the estimated stiffness from the respective fitted model, with the shaded region being the 95% confidence intervals. Comparisons were made at efforts ranging from zero to 30% MVC, the highest target provided to participants.

## Discussion

The ankle, particularly the stiffness of the ankle, plays a critical role in maintaining balance in response to unexpected postural disturbances. Since ankle stiffness is highly sensitive to the stiffness of the Achilles tendon (18), previously reported age-related decreases in Achilles tendon stiffness (4, 27, 44) could readily decrease ankle stiffness. However, there are two main limitations of previous studies. First, nearly all measures of Achilles tendon stiffness have been made at higher loads (above 30% MVC), which are less relevant to everyday tasks. Second, many comparisons between younger and older adults are made at matched efforts, where age-related differences in strength present a confounding factor. This study sought to determine if there are age-related changes in ankle stiffness when accounting for strength-related differences, and to investigate the potential causes. We quantified ankle, muscle, and tendon stiffness simultaneously in younger and older adults from 0 to 30% MVC, which are lower activations than previously explored. Contrary to our hypothesis that older adults would exhibit a decrease in ankle stiffness, we found no age-related differences in ankle stiffness at matched loads below 30% MVC. Similarly, the stiffness of the triceps surae and Achilles tendon did not vary with age when compared at matched loads. We did observe differences in ankle, muscle, and tendon stiffness at matched efforts, due to older adults being significantly weaker than younger adults. Our results suggest that there are no age-related changes in the mechanical properties of the triceps surae and Achilles tendon when tested at matched loads, at least at the lower loads relevant to posture and balance that were tested in this work. Instead, previously reported differences with age are likely dominated by age-dependent differences in strength.

### Effect of aging on ankle, muscle, and tendon stiffness

At matched levels of musculotendon force, we did not observe an age-related decrease in Achilles tendon stiffness for the range of loads tested in this work. Nearly all previous measurements were made at higher loads (typically above 30% MVC) that coincide with the linear region of the tendon stress-strain curve (4, 27, 44). In contrast, our measurements are made at or below 30% MVC, within the non-linear toe-region (17). It is possible that there are age-related decreases in tendon properties at higher loads rather than within the toe-region. To test this possibility, we compared the estimated parameter *K_Tmax_* in Eq. 2 between older and younger adults. *K_Tmax_* represents the maximum, constant tendon stiffness at high loads observed in other studies (e.g., the stiffness of the linear region of the stress-strain curve) (4, 23, 27, 44). The estimated maximum tendon stiffness was 16% lower in older adults compared to younger adults (younger: 108 ± 4 N/mm; older: 91 ± 2 N/mm; mean ± standard deviation; p<0.001). Future work should investigate how aging impacts tendon stiffness along the entirety of the stress-strain relationship.

Similar to previous results, we did observe a significant decrease in Achilles tendon stiffness at matched efforts (4, 27, 44). Stenroth, et al. (44), who estimated tendon stiffness as the slope of the force-strain relation between 10 – 80% MVC, reported a 17% decrease in Achilles tendon stiffness in older adults, which is a similar magnitude to what we observed at 30% MVC. Our findings clarify that these differences are due to age-related changes in strength, not mechanical properties of the tendon. Together, these results emphasize the importance of assessing tendon stiffness at matched loads when knowledge about mechanical properties is desired.

Our estimates of muscle stiffness include both a passive and active stiffness component. Passive muscle stiffness primarily originates from the connective tissue within muscle and titin (13, 36, 38, 43). There are well-established age-related changes in muscle that primarily affect its passive properties, such as a progressive decrease in muscle size, an increase in connective tissue, and an increase in fatty infiltration (5, 32, 44). Assuming that these changes were present in our population, their net effect does not appear to impact muscle stiffness when load is held constant (Fig 5D). Examining how passive muscle stiffness changes as it is loaded was outside the scope of this study and requires future investigation.

In contrast to passive muscle stiffness, there is limited evidence that active muscle stiffness would be impacted by aging. We have demonstrated previously that our estimates of muscle stiffness are consistent with measures of muscle short-range stiffness (17). Short-range stiffness scales with the activation-dependent force within a muscle (6), and is thought to arise from the number of attached cross-bridges (31, 37). While aging has been associated with changes in cross-bridge kinetics (30), to our knowledge, there is no evidence that these changes alter muscle short-range stiffness. Thus, at matched forces, it was unsurprising that muscle stiffness was similar between younger and older adults.

Older adults’ decrease in ankle stiffness at matched efforts compared to younger adults could negatively impact their balance control. During standing, older adults could either exert the same effort as younger adults, resulting in lower ankle stiffness, or could exert more effort to attain similar stiffness. The first strategy would lead to reduced postural stability and the latter to increased fatigue. Neither approach would be beneficial for the control of balance. Future work is required to determine which of these strategies is adopted during postural control.

### Co-contraction in older adults

Some of our older adults co-contracted during the experiment. Interestingly, those who did also scored significantly higher on the CDC fall risk assessment (Table 1). This is in accordance with previous findings that older adults with higher co-contraction during a balancing task were at a greater risk of falling (34). Additionally, all older adults could adequately control the feedback (both the plantarflexion torque and tibialis anterior EMG) when perturbations were not present. However, when the PRBS perturbations were present, some older adults were unable to prevent tibialis anterior activation. Numerous factors could be related to their inability to prevent tibialis anterior activation during the perturbations (21, 33, 34), but identifying those relevant to our protocol was not the purpose of this study.

### Limitations

Our system identification approach for estimating muscle and tendon stiffness assumes that all plantarflexion torque is transmitted through the Achilles tendon and triceps surae (17). This not only omits contributions from other structures spanning the joint but also required us to eliminate trials in which there was substantial co-contraction, which occurred in nearly 30% of our older subjects. This assumption can also lead to small errors during passive conditions (17), though those would have negligible impact on our main findings, which were drawn from comparisons at higher levels of volitional torque.

## Conclusion

We sought to determine how aging alters ankle, triceps surae, and Achilles tendon stiffness. Contrary to our primary hypothesis, we did not observe significant differences in these measures between younger and older adults when compared at matched loads. We did observe differences at matched efforts due to differences in strength between our groups. These results indicate that comparisons should be made at matched loads if the goal is to evaluate age-related changes in ankle, muscle, or tendon mechanical properties. Additionally, age-dependent changes in the maximum ankle stiffness that can be achieved may contribute to older adults’ increased risk of falling. Future work is warranted to quantify ankle stiffness in younger and older adults during standing, as it may aid in developing targeted interventions to improve balance in older adults.

## Funding

Research reported in this publication was supported by the National Institute On Aging of the National Institutes of Health under Award Number F31AG069412. The content is solely the responsibility of the authors and does not necessarily represent the official views of the National Institutes of Health. Research supported by the American Society of Biomechanics’ Graduate Student Grant-in-aid.

## Author Contributions

KLJ, DL, SSML, and EJP conceived of the study and designed the experimental protocol and analyses. KLJ carried out the experiments, analyzed the data, and drafted the manuscript. KLJ, DL, SSML, and EJP edited the manuscript. All authors approved the final version.

## Conflict of Interest

The authors declare no competing interests.

## Data Availability

Data will be made available upon reasonable request.

## Notes

### Competing Interest Statement

The authors have declared no competing interest.

## Reference

1. Baudry S. Aging changes the contribution of spinal and corticospinal pathways to control balance. Exercise and sport sciences reviews 44: 104–109, 2016.

2. Besomi M, Hodges PW, Clancy EA, Van Dieën J, Hug F, Lowery M, Merletti R, Søgaard K, Wrigley T, and Besier T. Consensus for experimental design in electromyography (CEDE) project: Amplitude normalization matrix. Journal of Electromyography and Kinesiology 53: 102438, 2020.

3. Clarke EC, Martin JH, d’Entremont AG, Pandy MG, Wilson DR, and Herbert RD. A non-invasive, 3D, dynamic MRI method for measuring muscle moment arms in vivo: demonstration in the human ankle joint and Achilles tendon. Med Eng Phys 37: 93-99, 2015.

4. Coombes BK, Tucker K, Hug F, and Dick TJ. Age-related differences in gastrocnemii muscles and Achilles tendon mechanical properties in vivo. Journal of Biomechanics 110067, 2020.

5. Csapo R, Malis V, Sinha U, Du J, and Sinha S. Age-associated differences in triceps surae muscle composition and strength–an MRI-based cross-sectional comparison of contractile, adipose and connective tissue. BMC musculoskeletal disorders 15: 209, 2014.

6. Cui L, Perreault EJ, Maas H, and Sandercock TG. Modeling short-range stiffness of feline lower hindlimb muscles. Journal of biomechanics 41: 1945–1952, 2008.

7. Delabastita T, Bogaerts S, and Vanwanseele B. Age-related changes in Achilles tendon stiffness and impact on functional activities: a systematic review and meta-analysis. Journal of Aging and Physical Activity 27: 116–127, 2018.

8. Ebrahimi A, Loegering IF, Martin JA, Pomeroy RL, Roth JD, and Thelen DG. Achilles tendon loading is lower in older adults than young adults across a broad range of walking speeds. Experimental gerontology 137: 110966, 2020.

9. Elias LJ, Bryden MP, and Bulman-Fleming MB. Footedness is a better predictor than is handedness of emotional lateralization. Neuropsychologia 36: 37–43, 1998.

10. Epro G, McCrum C, Mierau A, Leyendecker M, Brüggemann G-P, and Karamanidis K. Effects of triceps surae muscle strength and tendon stiffness on the reactive dynamic stability and adaptability of older female adults during perturbed walking. Journal of Applied Physiology 124: 1541–1549, 2018.

11. Finley JM, Dhaher YY, and Perreault EJ. Contributions of feed-forward and feedback strategies at the human ankle during control of unstable loads. Experimental brain research 217: 53–66, 2012.

12. Gill TM, Murphy TE, Gahbauer EA, and Allore HG. Association of injurious falls with disability outcomes and nursing home admissions in community-living older persons. American journal of epidemiology 178: 418–425, 2013.

13. Gillies AR, and Lieber RL. Structure and function of the skeletal muscle extracellular matrix. Muscle & nerve 44: 318–331, 2011.

14. Hill AV. The heat of shortening and the dynamic constants of muscle. Proceedings of the Royal Society of London Series B-Biological Sciences 126: 136–195, 1938.

15. Horak FB, and Macpherson JM. Postural orientation and equilibrium. Handbook of physiology 1: 255–292, 1996.

16. Jacobs JV, and Horak FB. Cortical control of postural responses. J Neural Transm (Vienna) 114: 1339–1348, 2007.

17. Jakubowski KL, Ludvig D, Bujnowski D, Lee SS, and Perreault EJ. Simultaneous quantification of ankle, muscle, and tendon impedance in humans. IEEE Transactions on Biomedical Engineering 2022.

18. Jakubowski KL, Ludvig D, Perreault EJ, and Lee SS. Non-linear properties of the Achilles tendon determine ankle impedance over a broad range of activations in humans. Journal of Experimental Biology jeb. 244863, 2023.

19. Kearney RE, and Hunter IW. System identification of human joint dynamics. Critical reviews in biomedical engineering 18: 55–87, 1990.

20. Kuo AD. The relative roles of feedforward and feedback in the control of rhythmic movements. Motor control 6: 129–145, 2002.

21. Laughton CA, Slavin M, Katdare K, Nolan L, Bean JF, Kerrigan DC, Phillips E, Lipsitz LA, and Collins JJ. Aging, muscle activity, and balance control: physiologic changes associated with balance impairment. Gait & posture 18: 101–108, 2003.

22. Le Mouel C, and Brette R. Anticipatory coadaptation of ankle stiffness and sensorimotor gain for standing balance. PLoS computational biology 15: e1007463–e1007463, 2019.

23. Lichtwark G, and Wilson A. Optimal muscle fascicle length and tendon stiffness for maximising gastrocnemius efficiency during human walking and running. Journal of theoretical biology 252: 662–673, 2008.

24. Ludvig D, Whitmore MW, and Perreault EJ. Leveraging joint mechanics simplifies the neural control of movement. Frontiers in Integrative Neuroscience 16: 802608, 2022.

25. Luke SG. Evaluating significance in linear mixed-effects models in R. Behavior research methods 49: 1494–1502, 2017.

26. Mackey DC, and Robinovitch SN. Mechanisms underlying age-related differences in ability to recover balance with the ankle strategy. Gait & posture 23: 59–68, 2006.

27. Mademli L, and Arampatzis A. Mechanical and morphological properties of the triceps surae muscle–tendon unit in old and young adults and their interaction with a submaximal fatiguing contraction. Journal of Electromyography and Kinesiology 18: 89–98, 2008.

28. Magnusson SP, Beyer N, Abrahamsen H, Aagaard P, Neergaard K, and Kjaer M. Increased cross-sectional area and reduced tensile stress of the Achilles tendon in elderly compared with young women. The Journals of Gerontology Series A: Biological Sciences and Medical Sciences 58: B123-B127, 2003.

29. Miguez D, Hodson-Tole EF, Loram I, and Harding PJ. A technical note on variable inter-frame interval as a cause of non-physiological experimental artefacts in ultrasound. R Soc Open Sci 4: 170245, 2017.

30. Miller MS, and Toth MJ. Myofilament protein alterations promote physical disability in aging and disease. Exercise and sport sciences reviews 41: 93, 2013.

31. Morgan DL. Separation of active and passive components of short-range stiffness of muscle. American Journal of Physiology-Cell Physiology 232: 45–49, 1977.

32. Morse CI, Thom JM, Reeves ND, Birch KM, and Narici MV. In vivo physiological cross-sectional area and specific force are reduced in the gastrocnemius of elderly men. J Appl Physiol 99: 1050–1055, 2005.

33. Nagai K, Yamada M, Uemura K, Yamada Y, Ichihashi N, and Tsuboyama T. Differences in muscle coactivation during postural control between healthy older and young adults. Archives of gerontology and geriatrics 53: 338–343, 2011.

34. Nelson-Wong E, Appell R, McKay M, Nawaz H, Roth J, Sigler R, Third J, and Walker M. Increased fall risk is associated with elevated co-contraction about the ankle during static balance challenges in older adults. European journal of applied physiology 112: 1379–1389, 2012.

35. Okoro CA, Hollis ND, Cyrus AC, and Griffin-Blake S. Prevalence of disabilities and health care access by disability status and type among adults—United States, 2016. Morbidity and Mortality Weekly Report 67: 882, 2018.

36. Prado LG, Makarenko I, Andresen C, Krüger M, Opitz CA, and Linke WA. Isoform diversity of giant proteins in relation to passive and active contractile properties of rabbit skeletal muscles. The Journal of general physiology 126: 461–480, 2005.

37. Rack PM, and Westbury DR. The short range stiffness of active mammalian muscle and its effect on mechanical properties. J Physiol 240: 331–350, 1974.

38. Reyna WE, Pichika R, Ludvig D, and Perreault EJ. Efficiency of skeletal muscle decellularization methods and their effects on the extracellular matrix. Journal of Biomechanics 110: 109961, 2020.

39. Rogers MW, and Mille ML. Timing paradox of stepping and falls in ageing: not so quick and quick (er) on the trigger. The Journal of physiology 594: 4537–4547, 2016.

40. Rubenstein LZ, Vivrette R, Harker JO, Stevens JA, and Kramer BJ. Validating an evidence-based, self-rated fall risk questionnaire (FRQ) for older adults. Journal of safety research 42: 493–499, 2011.

41. Runge C, Shupert C, Horak F, and Zajac F. Ankle and hip postural strategies defined by joint torques. Gait & posture 10: 161–170, 1999.

42. Sheehan FT. The 3D in vivo Achilles’ tendon moment arm, quantified during active muscle control and compared across sexes. J Biomech 45: 225–230, 2012.

43. Smith LR, Lee KS, Ward SR, Chambers HG, and Lieber RL. Hamstring contractures in children with spastic cerebral palsy result from a stiffer extracellular matrix and increased in vivo sarcomere length. The Journal of physiology 589: 2625–2639, 2011.

44. Stenroth L, Peltonen J, Cronin N, Sipilä S, and Finni T. Age-related differences in Achilles tendon properties and triceps surae muscle architecture in vivo. Journal of Applied Physiology 113: 1537–1544, 2012.

45. Stewart AL, MILLS KM, King AC, Haskell WL, Gillis D, and Ritter PL. CHAMPS physical activity questionnaire for older adults: outcomes for interventions. Medicine & Science in Sports & Exercise 33: 1126–1141, 2001.

46. Sturnieks DL, Menant J, Delbaere K, Vanrenterghem J, Rogers MW, Fitzpatrick RC, and Lord SR. Force-controlled balance perturbations associated with falls in older people: a prospective cohort study. PloS one 8: e70981, 2013.

47. Sturnieks DL, Menant J, Vanrenterghem J, Delbaere K, Fitzpatrick RC, and Lord SR. Sensorimotor and neuropsychological correlates of force perturbations that induce stepping in older adults. Gait & posture 36: 356–360, 2012.

48. Tankisi H, Burke D, Cui L, de Carvalho M, Kuwabara S, Nandedkar SD, Rutkove S, Stålberg E, van Putten MJ, and Fuglsang-Frederiksen A. Standards of instrumentation of EMG. Clinical neurophysiology 131: 243–258, 2020.

